# Latent-Based Imputation of Laboratory Measures from Electronic Health Records: Case for Complex Diseases

**DOI:** 10.1101/275743

**Authors:** V. Abedi, M.K. Shivakumar, P. Lu, R. Hontecillas, A. Leber, M. Ahuja, A.E. Ulloa, M.J. Shellenberger, J. Bassaganya-Riera

## Abstract

Imputation is a key step in Electronic Health Records-mining as it can significantly affect the conclusions derived from the downstream analysis. There are three main categories that explain the missingness in clinical settings–incompleteness, inconsistency, and inaccuracy–and these can capture a variety of situations: the patient did not seek treatment, the health care provider did not enter the information, etc. We used EHR data from patients diagnosed with Inflammatory Bowel Disease from Geisinger Health System to design a novel imputation that focuses on a complex phenotype. Our approach is based on latent-based analysis integrated with clustering to group patients based on their comorbidities before imputation. IBD is a chronic illness of unclear etiology and without a complete cure. We have taken advantage of the complexity of IBD to pre-process the EHR data of 10,498 IBD patients and show that imputation can be improved using shared latent comorbidities. The R code and sample simulated input data will be available at a future time.

## I. Introduction

Given the complexity and high-dimensionality of Electronic Health Records (EHR) the missingness and the need for imputation are an inevitable aspect in any study that attempts to use such data for downstream analysis. The EHR is not designed for research purposes, even though the breadth and depth of this information can be used to improve care at many levels, including developing predictive models of prognosis and designing *in silico* clinical trials [1-3]. Furthermore, the level and extent of the missing values in health care systems at large are not random. In fact, there are three main categories that explain the missingness in clinical settings [4, 5] – *incompleteness, inconsistency, and inaccuracy –* and these can capture a variety of situations, including the following: the patient could have been cared for outside of the health care system where the data are collected, the patient did not seek treatment, the health care provider did not enter the information, the patient has expired, the missing value was not needed for standard of care. Therefore, imputation can dramatically lead to biased results. Furthermore, excluding variables or patients with high-level of missingness can also introduce bias and reduce the scope of the study significantly. To address these challenges, different imputation techniques have been proposed, and each have their own advantages and limitations.

In a recent study, 12 different imputation techniques that were applied to laboratory measures from EHR were compared. In general, authors found that Multivariate Imputation by Chained Equations (MICE) and softImpute consistently imputed missing values with low error [6]; however, in that study, analysis was restricted to 28 most commonly available variables. In another study, authors assessed the different causes of missing data in the EHR data and identified these causes to be the source of unintentional bias [7]. Comparative analysis of three methods of imputation (a Singular Value Decomposition (SVD) based method (SVDimpute), weighted K-nearest neighbors (KNNimpute), and row average for DNA microarrays showed that in general KNN and SVD methods surpass the commonly accepted solutions of filling missing values with zeros or row average [8]. However, comparing imputation for clinical data with DNA microarray can be misleading. Missingness in DNA microarray is likely missing-at-random due to technical issues unlike missingness in EHR which is missing-not-at-random. In another study, fuzzy clustering was integrated with neural network to enhance the imputation process [9]. Furthermore, research has been done to evaluate imputation methods for non-normal data [10]. Using simulated data from a range of non-normal distributions and a level of missingness of 50% (missing completely at random or missing at random), it was found that the linearity between variables could be used to determine the need for transformation for non-normal variables. In the case of a linear relationship, transformation can introduce bias, while non-linear relationship between variables may require adequate transformation to accurately capture the non-linearity. Furthermore, many of the techniques are optimized for smaller levels of missingness (the most commonly available measurements), yet most clinical datasets (including the EHRs) have a significant percentage of missingness for many of their variables. To address this problem, machine learning methods have also been proposed [11]. In fact, the challenges of imputation for EHRs are unique, and if left unaddressed the utility of the data becomes limited [12]. Consequently, even though, for smaller targeted studies, it could be possible to integrate additional modalities, or perform analytical evaluation through chart review, to determine a likely cause of missingness; for larger studies this becomes infeasible. For examples, missingness level for many of very important variables (ex: inflammatory markers such as CRP, vitamin levels, or even A1C levels, a common biomarker for diabetes) can easily reach 50% or more in many realistic large datasets. Finally, given the complexity and the scale of the problem, in many studies, MICE [13] remains the method of choice. The MICE [13] algorithm is regression-based and assumes that the non-missing variables can be used for the predicting missing variables. However, this assumption does not always hold in EHR, yet, given the high-level of redundancy and presence of highly correlated variables in the EHR, imputation by MICE still performs relatively well for large clinical datasets. A comprehensive overview of handling missing data in EHR is presented in [12].

In this study, we have developed a novel imputation strategy based on latent-based analysis integrated with clustering to group patients based on their comorbidities before imputation. We apply this method on 10,498 patients diagnosed with Inflammatory Bowel Disease (IBD) and show its strength in analyzing laboratory measures with missingness levels up to 95%.

## II. Methodology

### Dataset

IBD is a complex disease with two main manifestations: Ulcerative Colitis (UC) and Crohn’s Disease (CD). Both UC and CD are highly significant public health problem in the U.S. and worldwide due to increasing incidence, morbidity, and mortality [14]. Moreover, IBD is a complex disease with no clear etiology or associated risk factors. Utilization of EHR can help facilitate better understanding this condition and continuous improvement in the design of treatment strategies for a more personalized care path.

### Data Extraction and Processing

We have identified an IBD cohort from the EHR of Geisinger Health System. Inclusion criteria of this cohort were based on extraction of patient population based on diagnosis recorded for patients under their visits, admissions and currently active problems listed under problem list, based on ICD9 and ICD10 codes for CD and UC (see Table 1). In order to confirm this diagnosis on patients, qualifying criteria included either two or more outpatient encounters, or one or more inpatient admissions, or an entry into problem list with an active flag checked. We have extracted clinical laboratory measurements for this cohort using the Logical Observation Identifiers Names and Codes (LOINC) system (see Table S1). For comorbidities, we extracted all the diagnosis for all the patients based on the ICD9 as well as ICD10 codes. Comorbidity data included details from visits, admissions, and problem list. We identified 10,498 patients with both comorbidity and laboratory data in the EHR. A total of 129 laboratory values, and 976 ICD codes (using 2-digit roll-up) were used. We excluded laboratory values with more than 95% missingness.

**Table 1:** ICD9 and ICD10 used for inclusion of Inflammatory Bowel Disease patients.

| Diagnosis | Inclusion Criteria using ICD codes |
| --- | --- |
| **ICD9 Diagnosis: Crohn's and Ulcerative Colitis** | 555.0, 555.1, 555.2, 555.9, 556.0, 556.1, 556.2, 556.3, 556.5, 556.6, 556.8, 556.9 |
| **ICD10 Diagnosis: Crohn's and Ulcerative Colitis** | K50.00, K50.011, K50.012, K50.013, K50.014, K50.018, K50.019, K50.10, K50.111, K50.112, K50.113, K50.114, K50.118, K50.119, K50.80, K50.811, K50.812, K50.813, K50.814, K50.818, K50.819, K50.90, K50.911, K50.912, K50.913, K50.914, K50.918, K50.919, K51.00, K51.011, K51.012, K51.014, K51.018, K51.019, K51.20, K51.211, K51.212, K51.213, K51.218, K51.219, K51.30, K51.311, K51.313, K51.314, K51.318, K51.319, K51.411, K51.414, K51.419, K51.50, K51.511, K51.513, K51.514, K51.518, K51.519, K51.80, K51.811, K51.812, K51.813, K51.814, K51.818, K51.819, K51.90, K51.911, K51.912, K51.913, K51.914, K51.918, K51.919 |

Two versions of the laboratory values were created. In the first version, the actual laboratory values (continuous values) were kept and the median values for each measurement were calculated. In the second version, all the laboratory values were converted to binary values using the flag in the system prior to calculating the median values. For instance, if a value was above or below the threshold (abnormal flag), then that value was converted to one; while for a value within the normal range, a zero was recorded (see Fig. 1). As a pre-processing step, any value that deviated more than 3 standard deviation from the mean was removed. Furthermore, a 0.1% white noise (error noise) was added to the binary values. Four different datasets were created, each with a limit on the level of missingness, ranging from low (5%), to moderation (50% and 75%), to extreme (95%).

**Fig. 1.**
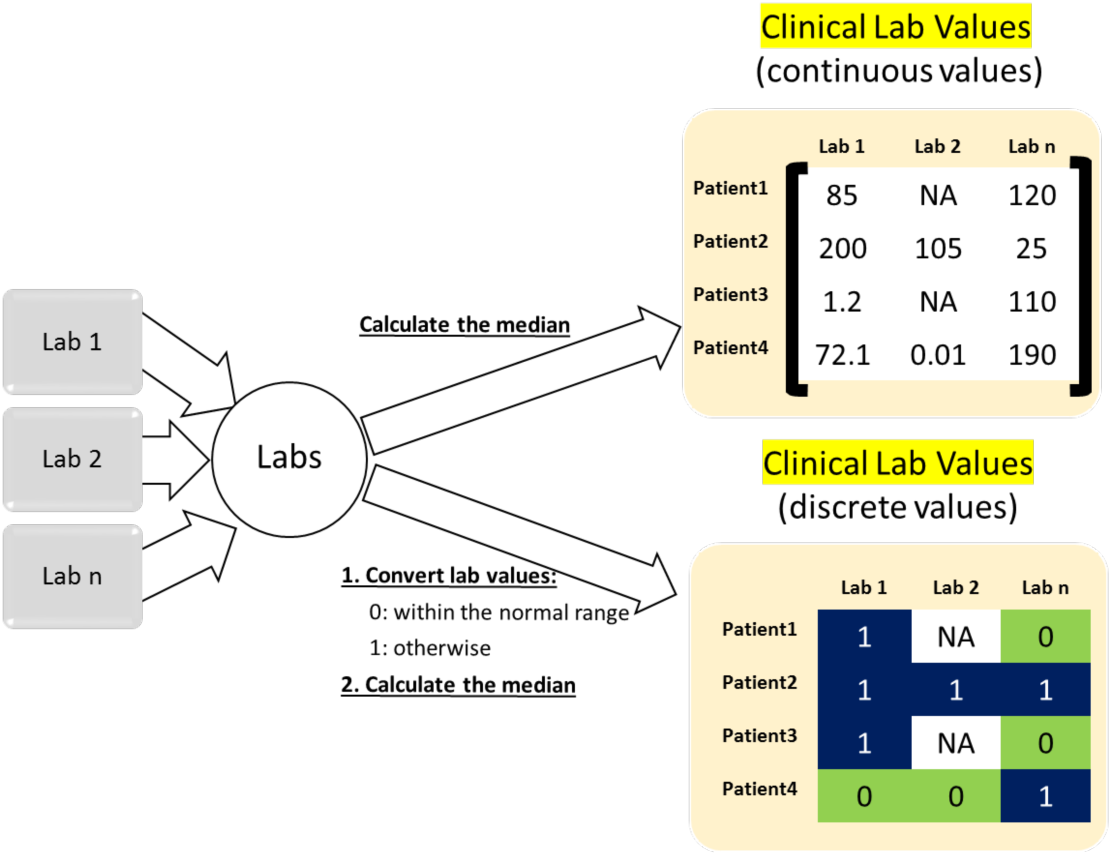
Construction of laboratory value matrices from EHR.

A binary comorbidity matrix for our cohort was also created using data from EHR. Data from primary or secondary diagnosis, as well as problem list were used for this purpose. During the first phase, ICD9/10 codes were compiled for each patient. During the second phase, the matrix was compressed using 3-digit roll up strategy, where ICD codes were combined if the first three digits were identical (ex: 289.52 and 289.53 --> 298). Furthermore, an ICD9/10 code was removed from the patient’s record if it was referenced only once. The resulting matrix was then converted to binary to represent the presence or absence of an ICD code for each patient. During the final stage, singular value decomposition (SVD) was applied to the matrix (Eq. 1) to compute the encoding of the dataset.

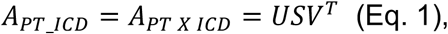

Where A_PT_ICD_ is the matrix encompassing all the ICD9/10 codes (presence of absence) for all the patients; U is an mxm matrix, S is an mxn diagonal matrix, and V is an nxn matrix. The colums of V are eigenvectors of A^T^A, and colums of U are eigenvectors of AA^T^. The diagonal elements of S are the square root of the eigenvalues of A^T^A or AA^T^.

The encoding matrix was then used to create different level of data abstraction by retaining only 10, 15, 66, or 85% of the encoding using dimensionality reduction technique (Eq. 2). Note that the approximation matrix is referred to as the data abstraction. The finalized output is referred to as latent comorbidities. Figure 2 summarize these steps.

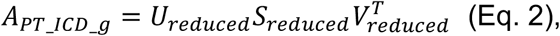

Where *g* is the level of abstraction (10%, 15%, 66%, and 85%), corresponding to the level of reduced matrices. A_PT_ICD_g_ is an approximation of the initial matrix (A_PT_ICD_).

**Fig. 2.**
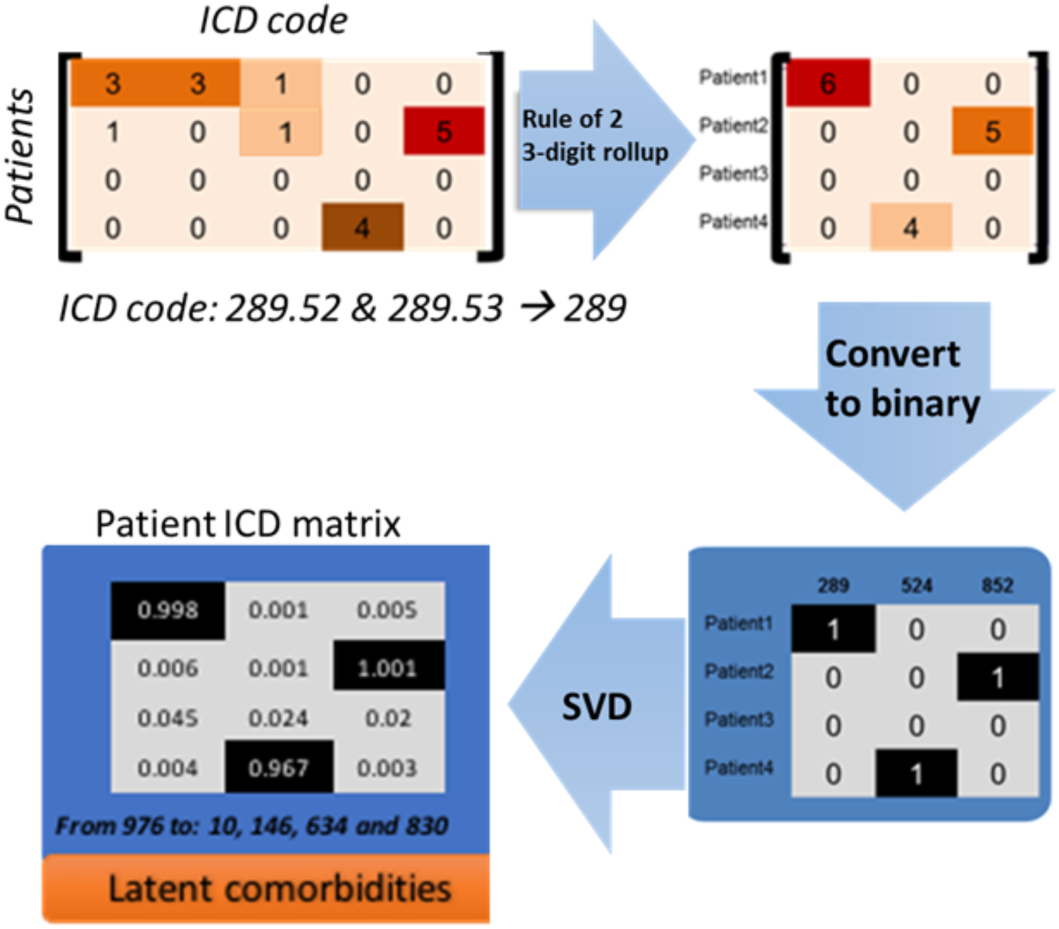
Construction of “Patient ICD Matrix” using comorbidity data from EHR.

### Imputation Strategies

#### Latent-based model

Our imputation method is a *hybrid* method, referred to as “latent-based model", that is based upon concurrently applying dimensionality reduction and clustering strategy. This hybrid method, efficiently captures relationships among features (or variables) and reduces noise (through dimensionality reduction) while providing an adaptive mechanism to perform imputation for any complex phenotype or trait. Using latent comorbidity data, patients were clustered using k-mean clustering technique. The data was clustered into 2, 4, 8, 16, 32 and 64 clusters. Imputation, using MICE, was then applied to each sub-group independently to predict the missing values. Furthermore, different variations in implementation were also explored.

#### Latent-based model with reduced sparsity

using the information from the singular value decomposition, the comorbidity data were evaluated for sparsity. The SVD captures the encoding of the matrix and insignificant ICD9/10 codes are transformed to values close to zero. Therefore, removing ICD9/10 codes corresponding to low values will reduce the sparsity of the matrix without eliminating important information. Three levels were assessed and number of disease codes (ICD9/10) was reduced to build three different datasets with varying level of sparsity. High sparsity is referred to when the sum of all the values for a ICD9/10 code in latent comorbidity matrix is less than 1 (that is the final matrix is very sparse by including the majority of the ICD9/10 codes), medium when the sum is less than 10 and low sparsity is when the sum is below 100 (only highly used ICD9/10 codes are included in the dataset). This strategy will also reduce noise, while also decreasing the size of the matrix, which will slightly reduce the computational complexity.

#### Feature expansion

a feature-expanded model was developed to evaluate if addition of comorbidity data as added features to the lab measurements could enhance the imputation performance as compared to the implementation of the hybrid model. In this situation, the matrix with the binary laboratory measures was expanded with the addition of comorbidity matrix (see Fig 3). The MICE imputation was then performed to predict the values of missing lab measurements.

**Fig. 3.**
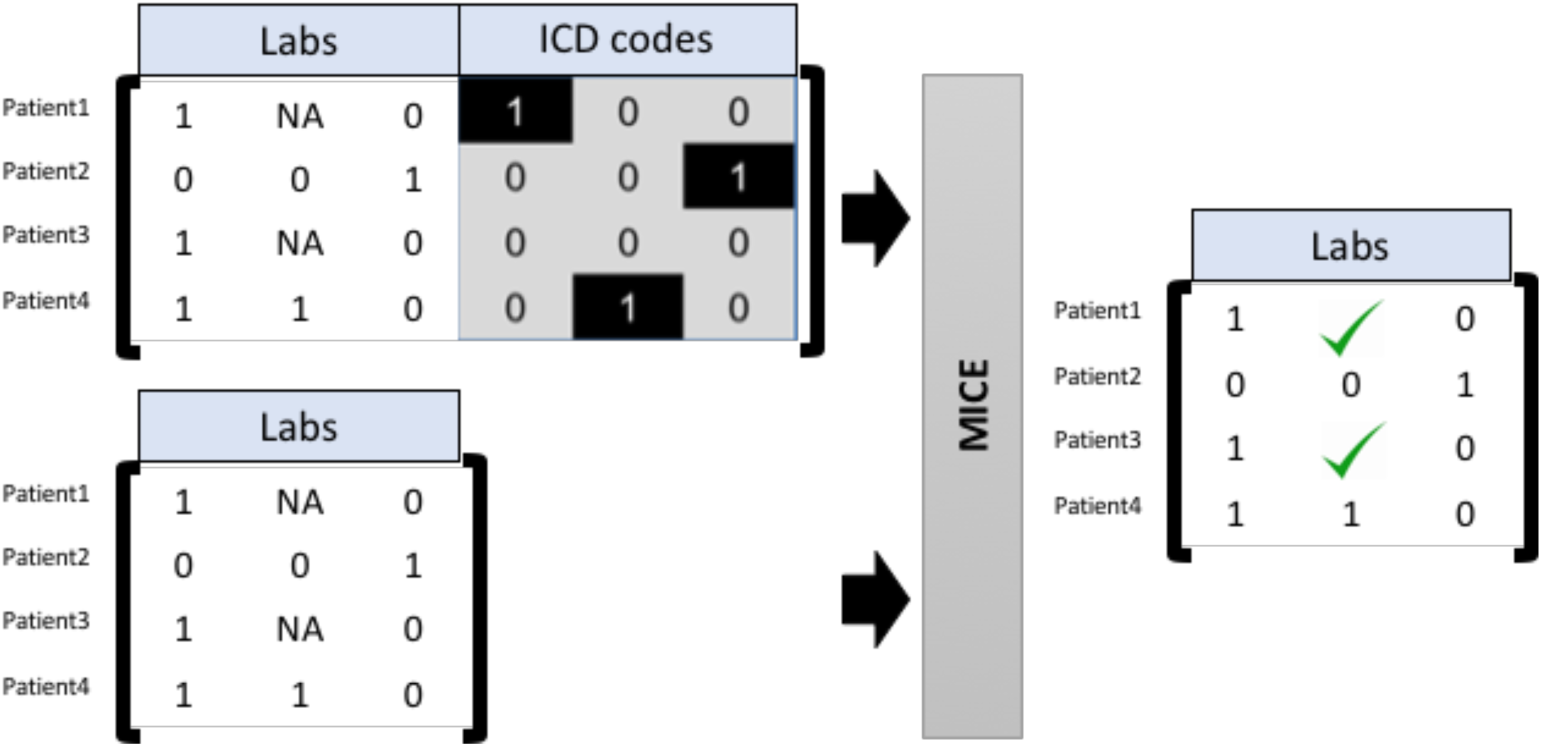
Feature expansion model. Laboratory and ICD codes are combined prior to imputation. The comparison is made with performing imputation on laboratory values using the same MICE algorithm.

#### Evaluation Strategy

Model evaluation was performed by randomly selecting variables and predicting them using the different strategies. To ensure fair distribution of random values that were withheld for testing before imputation, the data were split into bins based on missingness. The data were split into 2 bins in case of 5% missingness and 4 bins in case of 50%, 75% and 95% missingness. Equal number of values were withheld from each of the bins. Probability is low for selecting values from labs with high sparsity, thus sampling from bins ensures random values chosen to be withheld for testing are picked from all the levels of missingness. Each run was then repeated 10 times and the P-value was calculated using standard t-test statistics. The root mean square error (RMSE) was also calculated and averaged over the 10 runs. Comparison was based on calculating the difference between running imputation using the latent-based model and its’ various derivations techniques, and the standard MICE algorithm.

## III. Results

### A rich clinical dataset for Inflammatory Bowel Disease facilitates the development of imputation techniques targeted for complex phenotypes

We have identified 10,498 IBD patients from the Geisinger EHRs, with rich longitudinal data, spanning an average of nine years (time between first time diagnosed and current age) with over 10% of the IBD patients having over 15 years of EHR data in the system capturing their treatment experience and co-morbidities. Furthermore, only less than 20% of our IBD patients have less than five years of clinical data for analysis, making this dataset rich. In general, we observed, that for a very small set of the variables, the percentage of missingness was very small; however, the majority of the laboratory values had a relatively larger number of missing level. Fig. 4 highlights this trend. These observations further motivate the need for development of a method that is robust for imputation of sparse datasets.

**Fig. 4.**
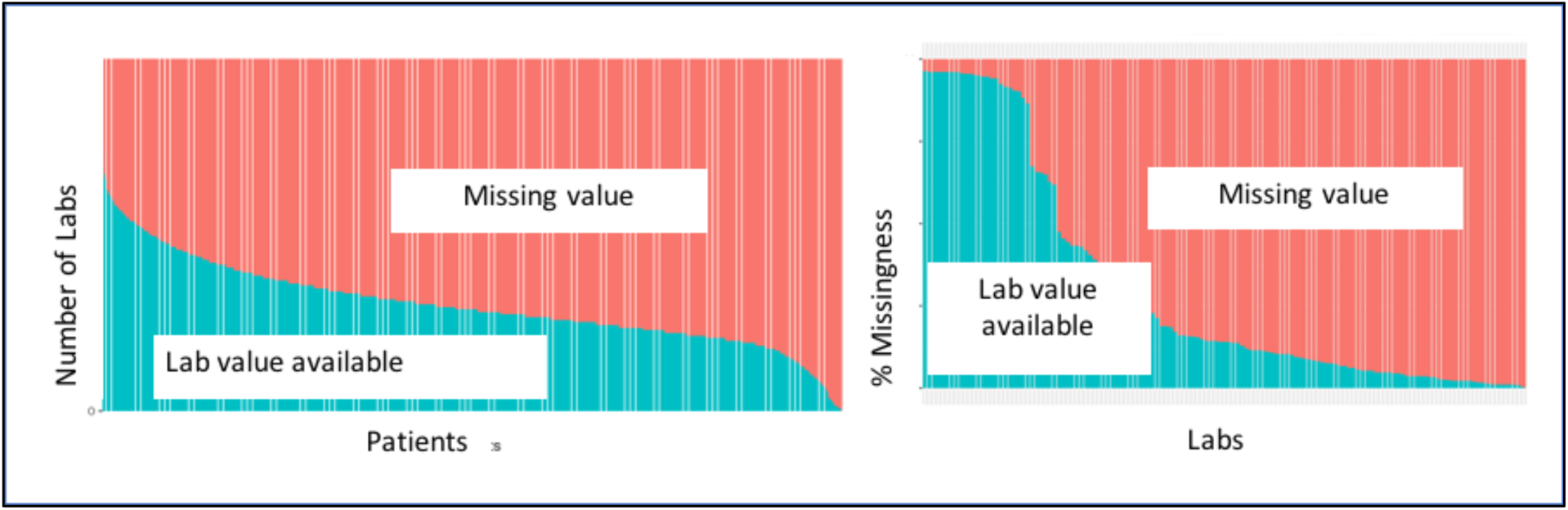
Missingness level in the Inflammatory Bowel Disease (IBD) cohort. Inflammation markers such as CRP (missing at 56%) and Sedimentation Rate (missing at 33%) are important lab measures for IBD.

### Comorbidity can *improve* imputation for complex phenotypes

Our novel method when compared with standard MICE algorithm shows significant improvement for variables with up to 95% missingness (Fig 5), even the comparison was disadvantageous for the hybrid method since the number of patients was larger when the data was not clustered. The goal of this study was to identify a combination of model parameters (number of clusters, data abstraction, and sparsity level), such that the imputation can be optimized for a given level of missingness in the data extracted from EHR. For instance, if low level of missingness is what is required, then for this specific dataset, optimal imputation can be performed by finding optimal region with that setting, considering both improved performance and significance level (P-value). However, if more lab measurements are of interest with possibly reaching a 75% missingness for some lab values, then panel A3,7,11 can be compared with their respective B3,7,11 P-value levels (see Fig. 5). Furthermore, if binary values are of interest as opposed to actual values, then one may consider evaluating sub-panels C1-12 and their corresponding p-values (D1-12). This strategy demonstrates the need to better understand the scope of the imputation technique prior to estimating the missing values from large and complex dataset retrieved from EHRs.

**Fig. 5.**
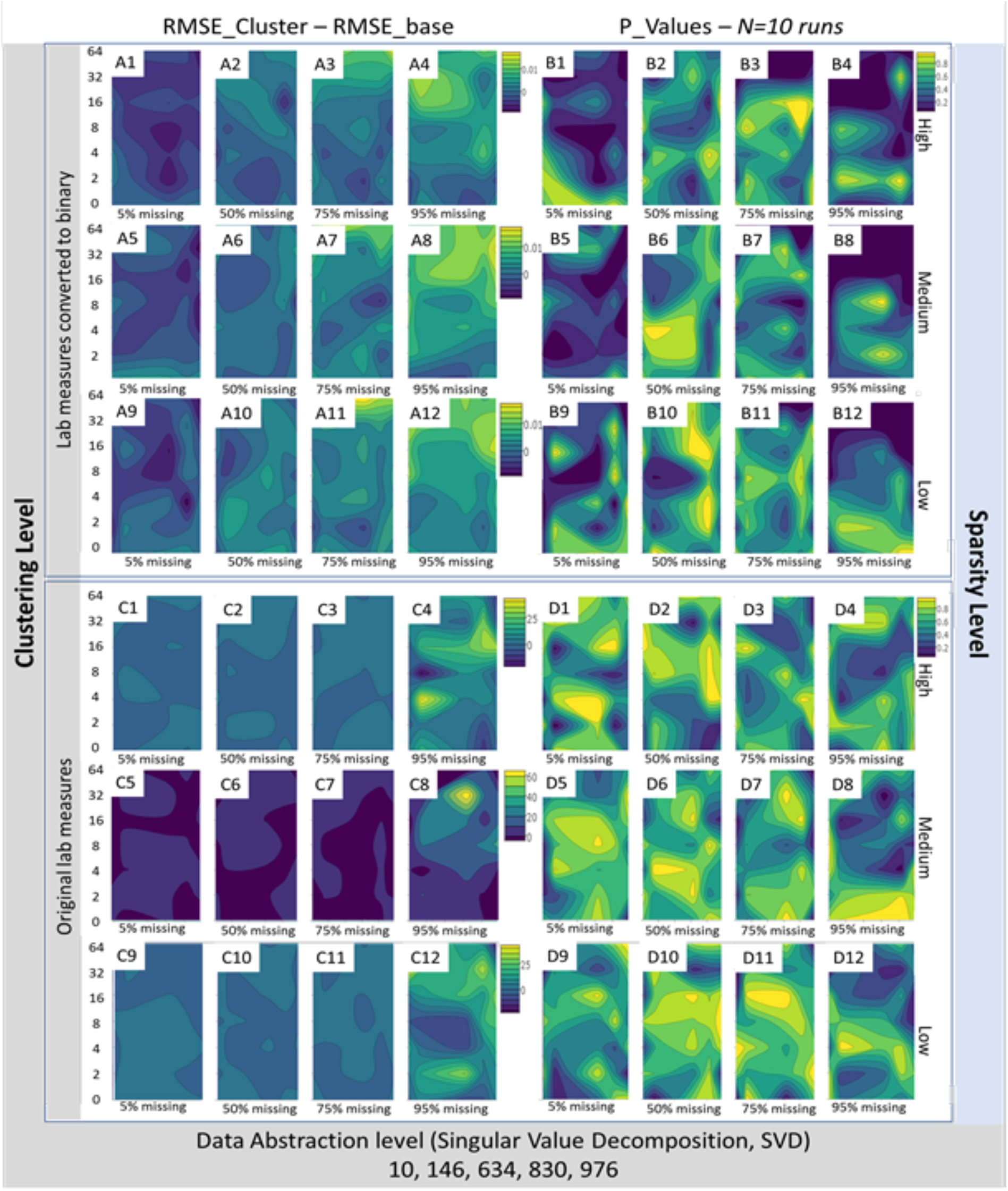
Optimized imputation method based on shared latent comorbidity information from EHR. The darker region corresponds to improvement of imputation using our novel method when compared to the standard MICE algorithm (difference between RMSE using new approached and MICE). Clustering level is based on number of clusters using k-mean clustering with *k* ranging from 0 to 64. Data abstraction level is based on using SVD and retaining only 10, 146, 634, 830 or 976 (or 10%, 15%, 66%, 85%, or 100%) of eigenvalues. Sparsity level is measured by including ICD codes that are prevalent using information from the encoding matrix from the SVD process; high sparsity (A1-4 and C1-4) is when all ICD codes are included in the model. Panels A1-12 correspond to imputation applied to binary lab measurement with their respective P-value (panels B1-12). Panels C1-12 correspond to imputation applied to original values with their respective P-value (panels D1-12).

### Increasing the feature space *deteriorates* the performance of imputation

By considering the feature expansion model alternative, we have increased the dimensionality of our feature space from 129 features (lab values) to about 1,000 features (129 labs + 976 ICD9 codes) for a fix population of 10K+ IBD patients. This alternative proved to be outperforming slightly the standard MICE, but for only the 5% missingness level (see Fig. 6). For higher level of missingness, the extended model did not outperform the standard MICE with the 129 variables.

**Fig. 6.**
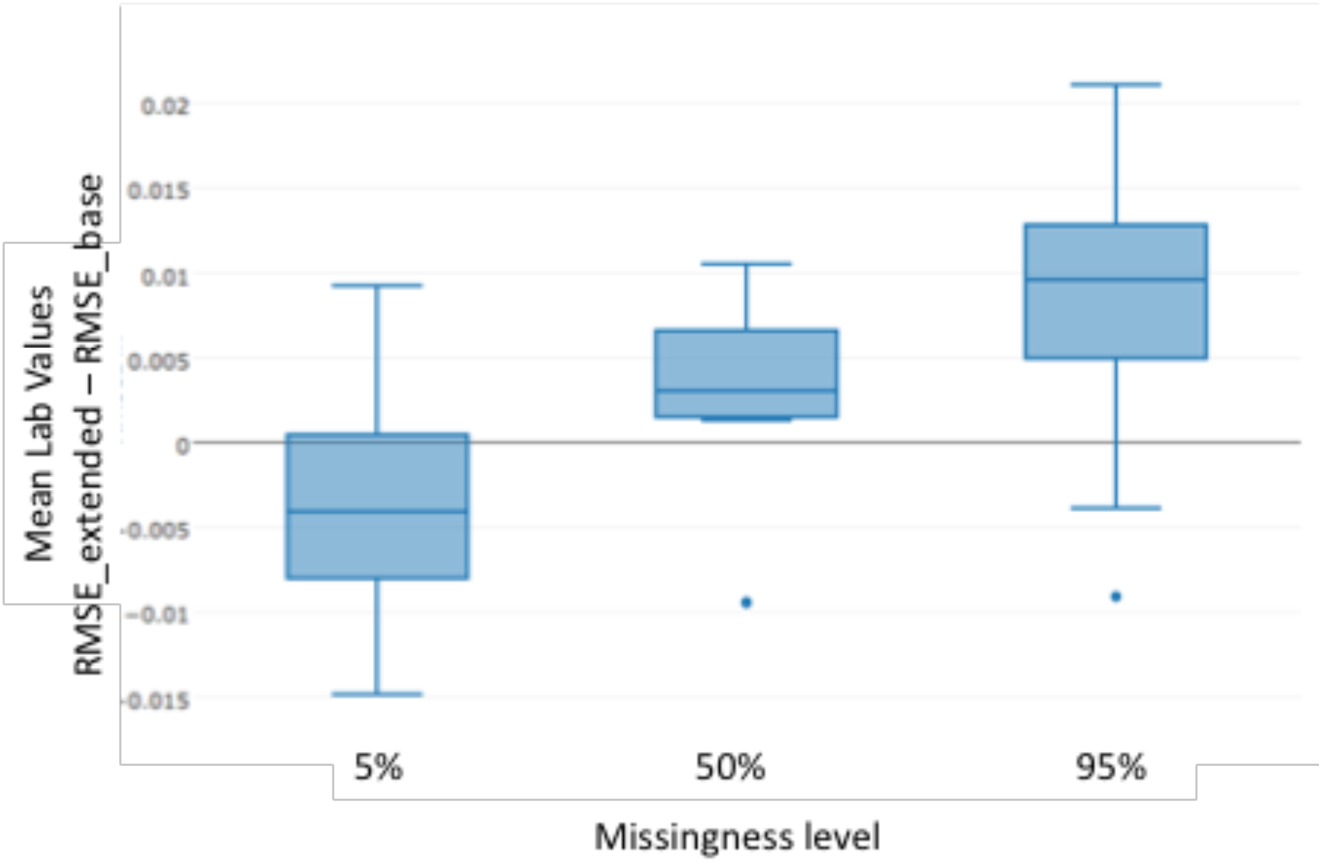
Increasing the feature space by using ICD code matrix information; N=20.

## IV. Discussion and Conclusion

The use of heterogeneous and large-scale clinical datasets, such as EHRs, provide an avenue for exploration of strategies to improve care at individualized levels, which include developing personalized models of response to therapy for complex diseases such as IBD. However, the data extracted from EHRs is noisy and have a large number of missing values, which requires a robust imputation strategy designed for large dataset of heterogeneous population. For realistic applications, It is not recommended to solely rely on redundancy of EHR data to conduct imputation, as our imputation strategy suggests. Our imputation strategy attempts to address this critical challenge using the IBD as a case study.

The IBD population is heterogeneous and a clear understanding of its risk factors is still lacking; therefore, treatment plan is usually still designed based on the average patient. Recent advances in knowledge of IBD pathogenesis have led to the implication of a complex interplay between metabolic reprogramming and immunity [15]. Furthermore, response to treatment in IBD varies significantly among individuals and disease subtypes based on demographic characteristics, diet, comorbidities, underlying immunological factors, and genetic polymorphisms. Thus, there is an urgent unmet clinical need to replace current approaches with personalized strategies that consider individual variability and diversity, and clinical data, if properly used, can provide a valuable resource in this field.

There has been a large number of studies developing novel imputation methods, and various studies assessing the use of different methods for large clinical studies. However, these methods do not tend to focus on a disease and more importantly on a complex disease or trait, rather they focus on the scale of the problem (number of cases, number of variables). It is only in some instances that studies are designed to investigate the possible correlation between variables and the effect of such correlation on the imputation outcome. However, majority of imputation methods for large clinical datasets (such as EHRs) will be used in a well-defined context, such as a specific disease or disease category (ex: infectious diseases, immune diseases, cancer, or cardiovascular diseases). In general, in a given study, the focus is rarely on all diseases at the same time, therefore, methods developed for imputation should also be designed with that in mind. Easy access to EHRs from large hospital systems could be one of the major limiting factors in the lack of development of novel imputation techniques for data from EHRs.

In this study, we have developed an intuitive method that can be applied for predicting missing values for complex diseases from EHRs. We have used the EHR s from Geisinger Health System, a pioneering institution in personalized and precision medicine. Geisinger has a national reputation for both high quality patient care and for being an innovative, integrated health care delivery system. This is especially true around electronic health information, being an early adopter of EPIC (1996) and building an enterprise-wide clinical data warehouse that contains comprehensive clinical and insurance claims data. We have reported demographic and clinical characteristics of the active patients in a recent publication [16].

The method presented here is an intuitive approach for any given complex disease, where bio-signatures or risk factors are only partially known and the relationship among the variables can be convoluted given the large dimensionality of the dataset. Furthermore, level of missingness can be high, even though the best results are typically obtained when the level of missingness is low or moderate. However, given the disease or condition of interest, the investigator will have the ability to include variables that may be crucial for the study. For instance, in the case of IBD, key variables of interest include CRP (missing at 56%), sedimentation rate (missing at 33%) which are not among the common variables in our dataset. Therefore, selecting the most commonly used variables will exclude important parameters for an investigation into the IBD cohort.

Using this approach, we were able to predict missingness of laboratory values that had missing level up to 95%. Our method was able to better predict the missing values than the standard algorithm. Furthermore, we have also shown that by simply integrating more features into the model and building very high-dimensional dataset (with about 1000 features) may not be an appropriate strategy, at least for a cohort of 10,000+ patients. Therefore, the hybrid-based strategy may be a better alternative in most situations.

Even though we had access to a large cohort of IBD population, our dataset was still limited if compared to more common conditions such as Diabetes with over 80,000 patients in our Geisinger cohort, or our COPD cohort of >49,000 patients. For those cases, more clusters could be formed to further improve the prediction outcome. Furthermore, we have only imputed median values; however, given a disease such as IBD, evaluating laboratory measures during the various stages of the disease may be more useful and also more predictive of the disease progression and relapse. However, imputing laboratory measures for each encounter is challenging at many levels, one such challenge is the significantly higher level of missingness in the dataset. As a future direction, we will investigate how to best impute longitudinal laboratory measures to better inform clinical studies. In addition, we will also explore integrating additional features, such as demographic information, age, gender, medication usage, as well as genetic information when available to further enhance the imputation outcome. Finally, we will evaluate various pre-processing and normalization strategies and evaluate if these manipulations can improve the outcome of our predictions, especially for variables with non-normal distributions. At last, even though MICE is one of the most popular imputation technique for EHRs, it would be interesting to compare our method with other algorithms.

To conclude, we have optimized the level of abstraction needed to improve the imputation for our IBD cohort of 10,498 patients. Our novel method when compared with standard “multiple imputation using chained equations” (MICE) shows significant improvement of up to 49% for variables with up to 95% missingness. We have shown that imputation can be improved, using *shared latent comorbidities*, and can facilitate generating a robust EHR dataset for further mining and modeling.

## Supplemental Material

**Table S1:** Laboratory data elements used in this study and their level of missingness in the IBD cohort.

| LAB LOINC Name | Missing % |
| --- | --- |
| Hemoglobin [Mass/volume] in Blood | 3.6 |
| Hematocrit [Volume Fraction] of Blood by Automated count | 3.9 |
| Leukocytes [# /volume] in Blood by Automated count | 3.9 |
| Erythrocyte mean corpuscular volume [Entitic volume] by Automated count | 3.9 |
| Erythrocytes [# /volume] in Blood by Automated count | 3.9 |
| Erythrocyte mean corpuscular hemoglobin [Entitic mass] by Automated count | 3.9 |
| Erythrocyte mean corpuscular hemoglobin concentration [Mass/volume] by Automated count | 3.9 |
| Erythrocyte distribution width [Ratio] by Automated count | 3.9 |
| Platelets [# /volume] in Blood by Automated count | 4.1 |
| Creatinine [Mass/volume] in Serum or Plasma | 4.4 |
| Platelet mean volume [Entitic volume] in Blood by Automated count | 4.6 |
| Potassium [Moles/volume] in Serum or Plasma | 4.7 |
| Urea nitrogen [Mass/volume] in Serum or Plasma | 5.0 |
| Chloride [Moles/volume] in Serum or Plasma | 5.3 |
| Carbon dioxide, total [Moles/volume] in Serum or Plasma | 5.3 |
| Calcium [Mass/volume] in Serum or Plasma | 5.8 |
| Glucose [Mass/volume] in Serum or Plasma | 6.0 |
| Sodium [Moles/volume] in Serum or Plasma | 7.4 |
| Bilirubin.total [Mass/volume] in Serum or Plasma | 8.3 |
| Protein [Mass/volume] in Serum or Plasma | 8.8 |
| Aspartate aminotransferase [Enzymatic activity/volume] in Serum or Plasma by With P-5'-P | 9.6 |
| Alanine aminotransferase [Enzymatic activity/volume] in Serum or Plasma by With P-5'-P | 9.7 |
| Anion gap 3 in Serum or Plasma | 11.8 |
| Neutrophils [# /volume] in Blood by Automated count | 13.5 |
| Sedimentation Rate | 32.7 |
| Triglyceride [Mass/volume] in Serum or Plasma | 34.1 |
| Bilirubin.direct [Mass/volume] in Serum or Plasma | 34.4 |
| Cholesterol [Mass/volume] in Serum or Plasma | 35.2 |
| Cholesterol in HDL [Mass/volume] in Serum or Plasma | 37.3 |
| Cholesterol in LDL [Mass/volume] in Serum or Plasma by calculation | 38.2 |
| Lipase [Enzymatic activity/volume] in Serum or Plasma | 54.4 |
| C reactive protein [Mass/volume] in Serum or Plasma | 55.5 |
| Iron [Mass/volume] in Serum or Plasma | 56.8 |
| Immature granulocytes [# /volume] in Blood by Automated count | 56.8 |
| Cobalamin (Vitamin B12) [Mass/volume] in Serum or Plasma | 57.1 |
| Immature granulocytes/100 leukocytes in Blood by Automated count | 58.0 |
| Ferritin [Mass/volume] in Serum or Plasma | 59.6 |
| Iron binding capacity. unsaturated [Mass/volume] in Serum or Plasma | 61.0 |
| Phosphate [Mass/volume] in Serum or Plasma | 64.6 |
| Magnesium [Mass/volume] in Serum or Plasma | 65.3 |
| Hemoglobin A1c/Hemoglobin.total in Blood by HPLC | 66.7 |
| Glucose [Mass/volume] in Blood by Automated test strip | 67.8 |
| Amylase [Enzymatic activity/volume] in Serum or Plasma | 68.8 |
| Creatine kinase [Enzymatic activity/volume] in Serum or Plasma | 69.4 |
| Folate [Mass/volume] in Serum or Plasma | 69.4 |
| Thyroxine (T4) free [Mass/volume] in Serum or Plasma | 70.7 |
| Neutrophils.band form/100 leukocytes in Blood by Manual count | 70.7 |
| Neutrophils.band form [#]/volume] in Blood by Manual count | 72.4 |
| Glucose mean value [Mass/volume] in Blood Estimated from glycated hemoglobin | 73.2 |
| Iron saturation [Mass Fraction] in Serum or Plasma | 73.3 |
| Cholesterol in LDL [Mass/volume] in Serum or Plasma by Direct assay | 77.3 |
| Creatine kinase.MB [Mass/volume] in Serum or Plasma | 78.7 |
| Prostate specific Ag [Mass/volume] in Serum or Plasma | 81.1 |
| Urate [Mass/volume] in Serum or Plasma | 81.2 |
| IgA [Mass/volume] in Serum or Plasma | 81.5 |
| Rheumatoid factor [Units/volume] in Serum or Plasma | 82.8 |
| Creatine kinase.MB/Creatine kinase.total in Serum or Plasma by Electrophoresis | 83.9 |
| Lactate dehydrogenase [Enzymatic activity/volume] in Serum or Plasma by Lactate to pyruvate reaction | 83.9 |
| Calcium.ionized [Moles/volume] in Serum or Plasma by Ion-selective membrane electrode (ISE) | 84.4 |
| Microalbumin [Mass/volume] in Urine | 84.4 |
| Microalbumin/Creatinine [Mass Ratio] in Urine | 84.5 |
| Gamma glutamyl transferase [Enzymatic activity/volume] in Serum or Plasma | 85.2 |
| Metamyelocytes/100 leukocytes in Blood by Manual count | 85.7 |
| Oxygen [Partial pressure] in Arterial blood | 85.7 |
| Carbon dioxide [Partial pressure] in Arterial blood | 85.8 |
| Fractional oxyhemoglobin in Arterial blood | 85.9 |
| Bicarbonate [Moles/volume] in Arterial blood | 86.0 |
| pH of Arterial blood | 86.0 |
| Metamyelocytes [#]/volume] in Blood by Manual count | 86.1 |
| Parathyrin.intact [Mass/volume] in Serum or Plasma | 86.3 |
| Triiodothyronine (T3) Free [Mass/volume] in Serum or Plasma | 87.2 |
| Oxygen saturation in Arterial blood | 88.0 |
| Beta globulin/Protein.total [Pure mass fraction] in Serum or Plasma by Electrophoresis | 88.4 |
| Gamma globulin/Protein.total [Pure mass fraction] in Serum or Plasma by Electrophoresis | 88.5 |
| Albumin/Protein.total [Pure mass fraction] in Serum or Plasma by Electrophoresis | 88.5 |
| Reticulocytes/100 erythrocytes in Blood by Automated count | 88.6 |
| Protein [Mass/volume] in Urine | 89.2 |
| Oxygen/Inspired gas setting [Volume Fraction] Ventilator | 89.5 |
| Myelocytes [#]/volume] in Blood by Manual count | 89.6 |
| Base deficit in Arterial blood | 89.7 |
| Chloride [Moles/volume] in Blood | 89.9 |
| Natriuretic peptide.B prohormone N-Terminal [Mass/volume] in Serum or Plasma | 90.6 |
| C reactive protein [Mass/volume] in Serum or Plasma by High sensitivity method | 90.9 |
| Carbon dioxide, total [Moles/volume] in Blood | 90.9 |
| Base excess in Arterial blood by calculation | 91.3 |
| Follitropin [Units/volume] in Serum or Plasma by 2nd IRP | 91.6 |
| Creatinine [Mass/volume] in Blood | 91.8 |
| Natriuretic peptide B [Mass/volume] in Serum or Plasma | 92.0 |
| Thyrotropin [Units/volume] in Serum or Plasma by Detection limit <= 0.005 mIU/L | 92.4 |
| Glucose [Mass/volume] in Blood | 92.5 |
| Cortisol [Mass/volume] in Serum or Plasma | 92.8 |
| IgG [Mass/volume] in Serum or Plasma | 93.2 |
| Testosterone [Mass/volume] in Serum or Plasma | 93.3 |
| Carboxyhemoglobin/Hemoglobin.total in Blood | 93.7 |
| IgM [Mass/volume] in Serum or Plasma | 93.8 |
| Methemoglobin/Hemoglobin.total in Blood | 94.4 |
| Lutropin [Units/volume] in Serum or Plasma | 94.6 |
| Cardiolipin IgG Ab [Units/volume] in Serum by Immunoassay | 94.8 |
| Cardiolipin IgM Ab [Units/volume] in Serum by Immunoassay | 94.8 |
| Carcinoembryonic Ag [Mass/volume] in Serum or Plasma | 95.2 |
| Testosterone Free [Mass/volume] in Serum or Plasma | 95.2 |
| Fibrinogen [Mass/volume] in Platelet poor plasma by Coagulation assay | 95.3 |
| Insulin [Units/volume] in Serum or Plasma | 95.3 |
| Thyroxine (T4) [Mass/volume] in Serum or Plasma | 95.4 |
| Homocysteine [Moles/volume] in Serum or Plasma | 95.5 |
| Calcidiol [Mass/volume] in Serum or Plasma | 95.9 |
| Albumin/Protein.total in Body fluid by Electrophoresis | 96.4 |
| Beta globulin/Protein.total in Serum or Plasma by Electrophoresis | 96.5 |
| Digoxin [Mass/volume] in Serum or Plasma | 96.6 |
| dRVVT (LA screen) | 97.2 |
| Alpha-1-Fetoprotein [Mass/volume] in Serum or Plasma | 97.4 |
| Methylmalonate [Moles/volume] in Serum or Plasma | 97.4 |
| Sex hormone binding globulin [Moles/volume] in Serum or Plasma | 97.7 |
| Cancer Ag 125 [Units/volume] in Serum or Plasma | 97.7 |
| Estradiol (E2) [Mass/volume] in Serum or Plasma | 97.8 |
| Testosterone.free+weakly bound [Mass/volume] in Serum or Plasma | 97.8 |
| Protein [Mass/time] in 24 hour Urine | 97.9 |
| Calcium [Mass/volume] in Urine | 98.0 |
| Calprotectin | 98.2 |
| Progesterone [Mass/volume] in Serum or Plasma | 98.3 |
| Prostate Specific Ag Free/Prostate specific Ag.total in Serum or Plasma | 98.7 |
| C peptide [Mass/volume] in Serum or Plasma | 98.8 |
| Glucose [Mass/volume] in Serum or Plasma --1 hour post dose glucose | 98.8 |
| Glucose [Mass/volume] in Serum or Plasma --2 hours post dose glucose | 98.9 |
| Aldosterone [Mass/volume] in Serum or Plasma | 98.9 |
| Glucose [Mass/volume] in Serum or Plasma --3 hours post dose glucose | 98.9 |
| Angiotensin converting enzyme [Enzymatic activity/volume] in Serum or Plasma | 99.0 |
| Volume of 24 hour Urine | 99.1 |

